# Alterations of Pleiotropic Neuropeptide-Receptor gene couples in Cetacea

**DOI:** 10.1101/2024.02.03.578746

**Authors:** Raul Valente, Miguel Cordeiro, Bernardo Pinto, André Machado, Filipe Alves, Isabel Sousa-Pinto, Raquel Ruivo, L. Filipe C. Castro

## Abstract

**Background:** Habitat transitions have considerable consequences in organism homeostasis, as they require the adjustment of several concurrent physiological compartments to maintain stability and adapt to a changing environment. Within the range of molecules with a crucial role in the regulation of different physiological processes, neuropeptides are key agents. Here, we examined the coding status of several neuropeptides and their receptors with pleiotropic activity in Cetacea.

**Results:** Analysis of 202 mammalian genomes, including 41 species of Cetacea, exposed an intricate mutational landscape compatible with gene sequence modification and loss. Specifically for Cetacea, in the twelve genes analysed we have determined patterns of loss ranging from species-specific disruptive mutations (e.g., Neuropeptide FF-Amide Peptide Precursor; *NPFF*) to complete erosion of the gene across the cetacean stem lineage (e.g., Somatostatin Receptor 4; *SSTR4*).

**Conclusions:** Impairment of some of these neuromodulators, may have contributed to the unique energetic metabolism, circadian rhythmicity and diving response displayed by this group of iconic mammals.

## Background

Evolutionary transitions, understood as the acquisition of a novel lifestyle within a lineage (e.g., [1]), encompass drastic modifications in several physiological systems (e.g., [2,3]), to sustain the impacts of a novel environment in physiological homeostasis. Cetacea, an iconic group of aquatic mammals composed of baleen whales (Mysticeti), toothed whales, dolphins and porpoises (Odontoceti), are a “*poster child for macroevolution*”, and typify such an evolutionary process [4]. The number of phenotypic modifications and adaptive traits associated with this land-to-water evolutionary transition is astonishing and implied a profound reorganization in different organ systems (e.g., [4, 5, 6]; Figure 1). In fact, many cetacean physio-anatomical traits set them apart as unique among mammals. One remarkable example is their circadian rhythmicity, which was adapted by abolishing neurochemical signals like the hormone melatonin and the neuropeptide cortistatin [5, 7, 8, 9]. This alteration was accompanied by anatomical and physiological rearrangements, as evidenced by their lower cortical orexinergic bouton density, when compared to closely related terrestrial artiodactyls [10]. Yet, the precise molecular events underpinning the coalescence of these interconnected systems into novel physiological states remain largely unexplored. Among the various chemical messengers modulating physiological functions, neuropeptides are central in regulating various physiological mechanisms [11]. They are defined as short sequences of amino acids synthesized and released by neurons or glia. Moreover, they are responsible for slow-onset, long-lasting modulation of synaptic transmission, acting on neighbouring cells via G protein-coupled receptors (GPCRs) [12].

**Figure 1:**
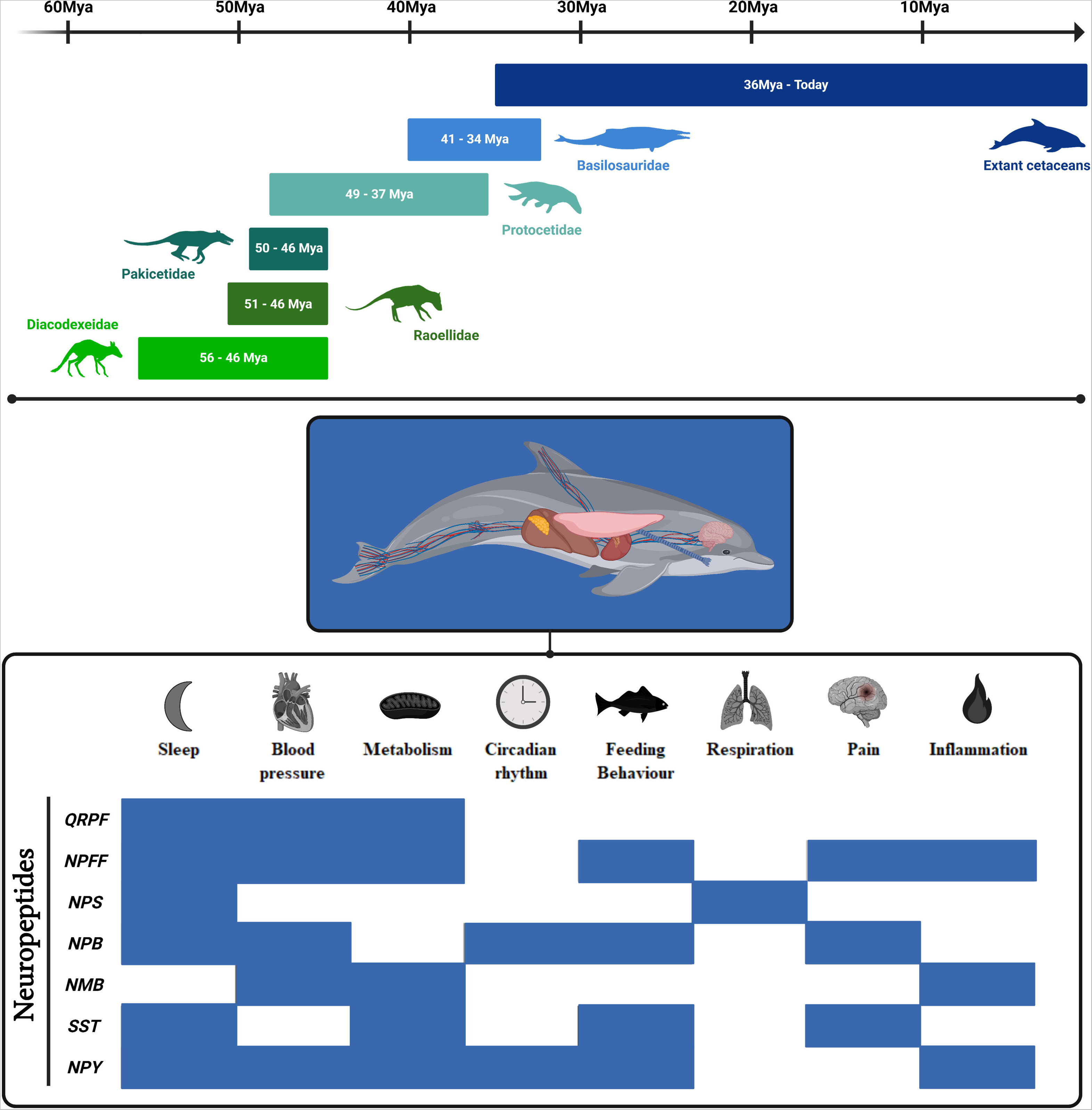
Evolutionary transitions and pleiotropic molecules. Land-to-water habitat transition experienced by marine mammals such as Cetacea was accompanied by remarkable anatomical, physiological, and behavioural adaptations. In this study, we aim to assess the coding status of several neuropeptides with distinct physiological roles. Fossil timeline (gray bars) depicts the estimated ages of the appearance and extinction of key cetacean ancestors. Dating of these events were obtained in the following references [100, 101, 102, 103, 104, 105, 106, 107]. In the lower panel, references for each reported role associated with neuropeptides is provided by [14, 18, 26, 60, 66, 69, 71, 73, 79, 108, 109, 110, 111, 112, 113, 114, 115, 116, 117, 118].

In this study, we delve into the evolution of several neuropeptide and receptors, with an important role in the coordination of different physiological compartments that have been clearly modified in Cetacea: including the liver, lungs, heart and brain (Figure 1; [13]). It is worth noting that neuropeptides and their receptors often exhibit pleiotropic effects, implying a potential wide range of physiological functions. As such, Neuropeptide FF-Amide Peptide Precursor (*NPFF*) seems a major player in pain modulation [14, 15], but also has a role in cardiovascular regulation (e.g., [16]). Pyroglutamylated RFamide Peptide (*QRFP*) and its receptor (*QRFPR*) play a role in the control of feeding behavior and also participate in sleep regulation [17, 18, 19]. Neuropeptide B (*NPB*) and one of its receptors (*NPBWR2*) are also important for the control of feeding behaviour, as well as pain modulation and sleep regulation [20, 21, 22, 23]. Neuropeptide S (*NPS*) and its receptor (*NPSR1*) are central to sleep regulation and are linked to the control of feeding behaviour [24, 25, 26]. Neuromedin B Receptor (*NMBR*) has been linked with both hormonal regulation and immunity control [27, 28]. Somatostatin Receptor 4 (*SSTR4*) plays a role in hormonal regulation and in stress responses [29]. Finally, Neuropeptide Y6 Receptor (*Npy6r*) was associated with hormonal regulation and control of feeding behaviour [30]. Such pleiotropy raises the critical question on how these regulators evolved in such a drastic ecological transition as that experienced in Cetacea evolution. Strikingly, by expanding the analysis to non-Cetacea 161 mammalian species (22 taxonomic orders; Additional Table 1), we also revealed a complex mutational landscape, possibly indicative of distinct adaptations to multiple environments.

## Material and Methods

### Genomic Sequence Compilation

An exhaustive literature review was used to identify a list of key genes that encode neuropeptides and respective receptors with pleiotropic action in physiological compartments that have suffered drastic adjustments in the land to water transition. Consequently, *NPFF*, *NPFFR1*, *NPFFR2*, *QRFP*, *QRFPR*, *NPS*, *NPSR1*, *NPB*, *NPBWR2*, *NMBR*, *SSTR4* and *Npy6r* corresponding genomic regions from 172 mammals with the genome annotated were collected from NCBI (Additional Table 1). Retrieval was performed using Genomic Sequence Downloader (https://github.com/luisqalves/genomic_sequence_downloader.py), using human synteny for each target gene as input – downstream and upstream flanking genes considered were the ones presenting a Gene type as “protein coding”. Whenever Genomic Sequence Downloader failed to obtain a genomic region containing the target gene, there was a tempt to collect directly the corresponding genomic sequences from the reference genome assemblies available at NCBI – either using the target gene or the corresponding flanking genes to tell us the genomic region comprising the physical location of the gene. In addition to the species previously referred, 29 genomes from cetaceans with no annotated genome available and *Hippopotamus amphibius* (hippopotamus) – either from NCBI, DNA Zoo [31] or Bowhead Whale Genome Resource (http://www.bowhead-whale.org) – were also inspected. In this case, the available genome assemblies were searched through blastn using *Bos taurus* target gene corresponding ortholog coding sequence (CDS), as well as the CDS’s of the flanking genes in the same species. Exception made for *SSTR4* where *H. sapiens* CDS for target gene and flanking genes were used as query. The best matching genome scaffold was retrieved. When no consensual blast hit was obtained, all hits corresponding to the query were inspected, the aligning regions submitted to a back-blast search against the nucleotide (nt) database of NCBI, with the matching genomic sequence(s) corresponding to the gene of interest being the one(s) selected for annotation (when existing).

### Inference of the Coding Status

Gene annotation was firstly performed using a pseudogene inference pipeline, Pseudo*Checker* [32]. For each run the *Homo sapiens* (human) orthologous gene and genomic sequences were used as reference (NCBI Accession ID regarding human *NPFF*: NM_003717.4; *NPFFR1*: NM_022146.5; *NPFFR2*: NM_004885.3; *QRPF*: NM_198180.3; *QRPFR*: NM_198179.3; *NPS*: NM_001030013.2; *NPSR1*: NM_207172.2; *NPB*: NM_148896.5; *NPBWR2*: NM_005286.4; NMBR: NM_002511.4; SSTR4: NM_001052.4; default parameters were maintained). For gene annotation in cetaceans and hippopotamus, *B. taurus* CDS were USED as reference, whenever it presented a curated transcript (NCBI Accession ID regarding cattle *NPFF*: NM_174123.3; *QRFP*: NM_198222.1; *QRFPR*: NM_001192681.1; *NPSR1*: NM_001192977.2; *NPB*: NM_173944.1; *NPBWR2*: NM_174075.1; *NMBR*: NM_001205710.1). Finally, for *Npy6r* annotation, *Mus musculus* (house mouse) orthologous gene was used as reference (NCBI Accession ID: NM_010935.4). Estimation of the erosion condition of the tested genes, was performed through PseudoIndex, a user assistant metric built into the software Pseudo*Checker*, that ranges on a discrete scale from 0 (coding) to 5 (pseudogenized) [32]. PseudoIndex considers three key components: absent exons, shifted codons and truncated sequences, each quantifying various aspects of mutational evidence. The predicted sequences were classified into “functional” if PseudoIndex was between 0 and 2, or “putatively pseudogenized” when PseudoIndex was higher than 2. Manual gene annotation and validation was performed for species presenting PseudoIndex higher than 2, with the collected genomic sequences being uploaded into Geneious Prime®2021.2.2 and the gene sequence manually predicted as described in Lopes-Marques [33] using the previous coding sequences from the same species as reference. Briefly, reference exons were mapped to the corresponding genomic sequences and subsequently aligned regions were manually inspected to find putatively ORF disrupting mutations (frameshifts (indels), premature stop codon, loss of canonical splice sites). Given the small size of the first exon in human *NPFFR1* and *NPS* (only 7 and 8 base pairs, respectively), putative inactivating mutations in these exons were not considered. The identified ORF disrupting mutations (one per species) were validated by searching at least two independent Sequence Read Archive (SRA) projects (when possible) of the corresponding species. Conserved deleterious mutations were validated in only one species.

### Assessment of Gene Expression in Multiple Tissues

The global analyses of relative gene expression in Cetacea species were performed in four cetacean species, using the approach described previously [34]. The genome and annotations of *D. leucas* (Accession number; GCF_002288925.2), *M. monoceros* (Accession number; GCF_005190385.1) and *B. acutorostrata* (Accession number; GCF_000493695.1) were downloaded from NCBI genome browser. On the other hand, the genome and annotation of *B. mysticetus* were retrieved from http://www.bowhead-whale.org/. Next, we collected the RNA-seq datasets from NCBI (Additional Table 2). When more than one dataset was available for a specific tissue, both were concatenated. Before proceeding with relative gene expression analyses, the genomes and annotations were checked for errors with (agat_convert_sp_gxf2gxf.pl) script and converted from .gff to .gtf format with (agat_convert_sp_gff2gtf.pl) script of AGAT tool [35]. The RNASeq datasets were mapped against each genome with the Hisat2 v.2.2.1 aligner [36, 37], and the relative gene expression determined with the StingTie v.2.2.1 software [38], following the authors protocol [39]. In the end, the gene expression quantifications were used in transcript per million (TPM).

To assess the quality of each transcriptomic dataset, we calculated gene expression in a highly expressed gene according to the Human Protein Atlas (http://www.proteinatlas.org). Genes included in this analysis were the following: Arginine Vasopressin (*AVP*; for brain), Phosducin (*PDC*; for retina); Uromodulin (UMOD; for kidney), Transition protein-1 (*TNP1*; for testes), Nicotinamide Riboside Kinase 2 (*NMRK2*; for muscle), Acrosin (*ACR*; for heart), Secreted Phosphoprotein 2 (*SPP2*; for liver), Advanced Glycosylation End-Product Specific Receptor (*AGER*; for lung), C-C Motif Chemokine Ligand 25 (*CCL25*; for lymph) and Leptin (*LEP*; for blubber).

To inspect the occurrence of expression reads from our target genes, mRNA reads previously collected were mapped to corresponding annotated genes using the “map to reference” tool available in Geneious Prime®2021.2.2 (maximum read gap size adjusted to the length of the maximum distance between exons with a maximum mismatch rate of 1%). The mapped reads were then classified as either spliced reads (spanning over two exons), exon-intron reads, or exonic reads, based on the genomic region they mapped to. After classification, the reads of each class were counted, and aligning regions covering mutated regions were screened for mutated transcripts.

### Molecular Evolutionary Analyses

Given the slow rate of evolution in Mysticeti [40], evidence for pseudogenization does not accumulate rapidly. For neuropeptide and receptor genes that undergo knockout in odontocetes, yet remain intact in mysticetes (*QRFP*, *NPS*, *NPSR1*, *NPB*, *NMBR*), estimation of the selection intensity acting on these genes (through dN/dS analyses) within Mysticeti would provide valuable insights. For this purpose, we translation-aligned the predicted sequences of Mysticeti (retrieved from Pseudo*Checker*) and several mammals from different lineages without any inactivating mutations (see Additional Table 3) using the Blosum62 substitution matrix in Geneious Prime® 2021.2.2. To investigate potential relaxation of purifying selection, often linked with gene pseudogenization, we conducted RELAX analysis using the HyPhy package [41, 42]. This analysis involves comparing a foreground set of species (Mysticeti in this case) with a background set (all non-cetacean species) within a hypothesis-testing framework. In cases where significant evidence of relaxation was found, we additionally run aBSREL [43] and BUSTED [44] to ascertain whether positive selection has manifested across a subset of branches or to assess gene-wide (rather than site-specific) positive selection. The idea behind these analyses is that if there is no evidence of positive selection, the accelerated rate in a particular clade probably indicates a pattern of relaxed purifying selection.

## Results

### Variable patterns of sequence alterations are found in two RFamide neuropeptides and their receptors

Comparative analysis of the *Neuropeptide FF-Amide Peptide Precursor* (*NPFF*) exonic sequences revealed localized events of gene pseudogenization across the mammalian tree. More specifically, 38 mammal species presented a PseudoIndex higher than 2 (Additional Table 4). This built-in assistant metric incorporated in Pseudo*Checker* [32], assess the sequence erosion status of the tested genes on a discrete scale ranging from 0 (coding) to 5 (pseudogenized).

Subsequent manual annotation, allowed the identification of non-conserved open reading frame (ORF) disruptive mutations in both Mysticeti and Odontoceti species, including loss of start codon or indels in exons 1 and 2 (Additional Table 5). When expanding our search to other mammalian lineages, deleterious mutations were found in kangaroo rats (*Dipodomys* spp.), Hawaian monk seal (*Neomonachus schauinslandi*) and flying foxes (*Pteropus* spp.) (Figure 3); in the latter a conserved mutation was validated by SRA searches (Additional File 1). Also, although no inactivating mutations were discovered, exons 1 and 2 in Canidae (*Canis* spp. and *Vulpes* spp.) were not found (Additional Table 5).

Next, we sought to characterize the coding status of both *NPFF* receptors. Pseudo*Checker* analyses for *Neuropeptide FF Receptor 1* (*NPFFR1*) resulted in 127 species presenting a PseudoIndex > 2, contrasting with *Neuropeptide FF Receptor 2* (*NPFFR2*) where only 48 species seem to yield eroded genes (Additional Tables 6 and 8). Annotation of collected cetacean genomic sequences revealed extension of the *NPFFR1* gene across all analysed species, due to a conserved 11 nucleotide deletion in exon 4 (Figure 2; Additional Table 7). For *NPFFR2*, most of the cetaceans displayed several disrupting mutations including a conserved 2 nucleotide deletion in exon 3 within Delphinoidea (oceanic dolphins, porpoises, beluga and narwhal) and within Physeteroidea (sperm whales) (Figure 2; Additional Table 9). In both these receptors the majority of the ORF-disrupting mutations are found in the last exon. which translates in ~20% of the total gene content completely modified (for *NPFFR1*) and for *NPFFR2* more than 25% of the total gene length is not transcribed given the change of frame due to a frameshift mutation.

**Figure 2:**
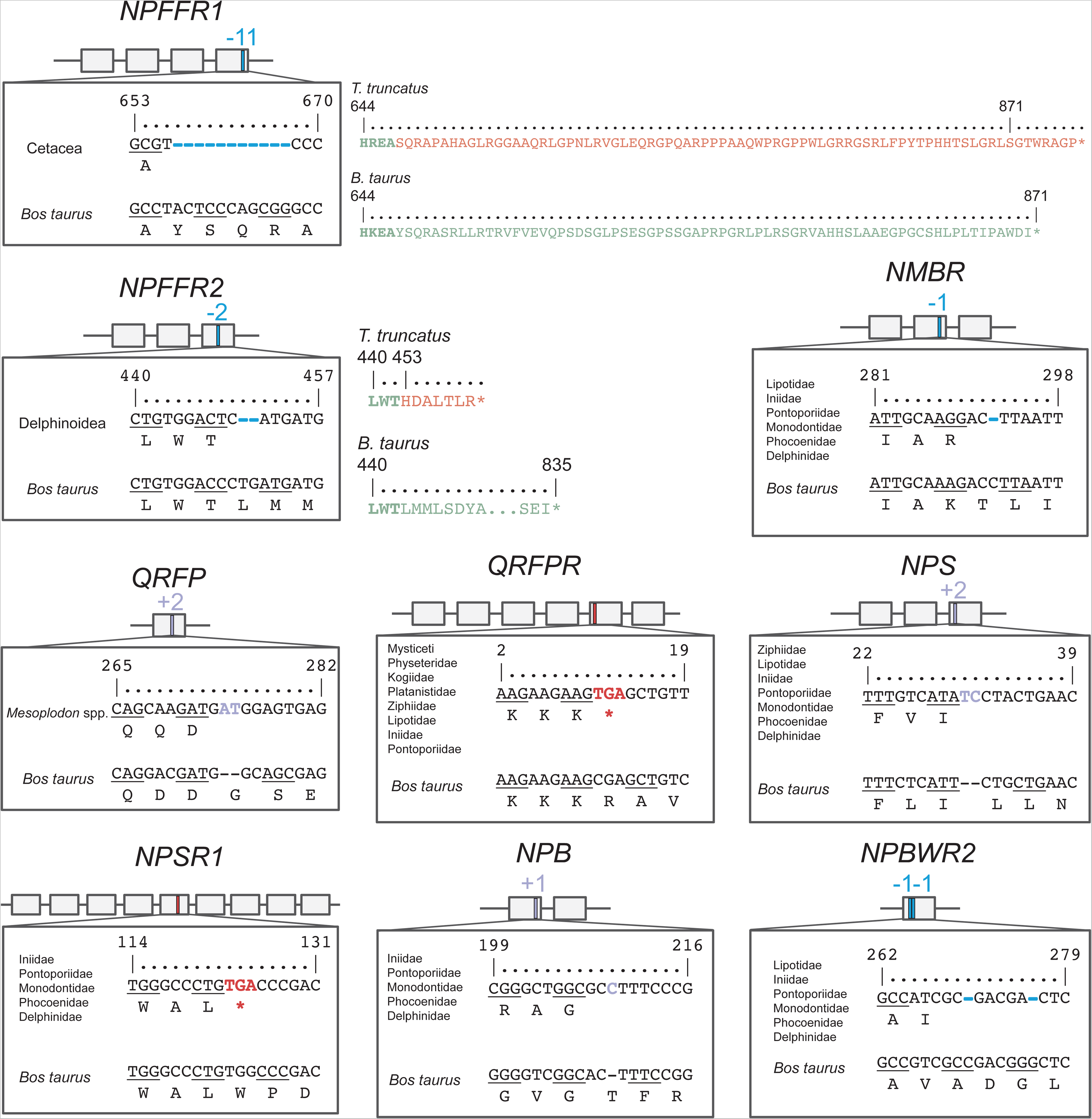
Pseudogene annotation in Cetacea. Examples of ORF-disrupting mutations in Cetacea for the study genes and respective sequence alignment. Each box represents an exon and lines represent intronic regions.

In other mammalian species, a highly eroded *NPFFR1* was also found within Chiroptera (bats), where the majority of the analysed species exhibited several conserved ORF-abolishing mutations and/or lack of exons (Figure 3; Additional Table 7). Moreover, ORF-disrupting mutations were also identified in other mammals, however data from independent SRAs only validated mutations in *Canis* spp., red fox (*Vulpes vulpes*) and (naked mole-rat (*Heterocephalus glaber*) (Additional Table 9). SRA validations for *NPFFR1* and *NPFFR2* are presented in Additional Files 2 and 3. Within Cetacea, Pseudo*Checker* analyses for *Pyroglutamylated RFamide Peptide* (*QRFP*) resulted in 20 species presenting a PseudoIndex higher than 2 (Additional Table 10). In cetaceans, most of the species showed either an eroded or sequence-modified *QRFP* with conserved mutations within *Mesoplodon* spp. (beaked whales) and Delphinidae (Figure 2; Additional Table 11). Specifically for baleen whales and oceanic dolphins presenting only mutations at the beginning of the gene, we may not rule out the possibility of having a functional *QRPF* in these species, as the peptide does not suffer any change of frame while transcribed. Investigation of *QRFP* inactivation in other mammalian lineages revealed episodes of gene loss (Figure 3). In particular, *QRFP* was found to be inactivated and subsequently validated in species such as *Pteropus* spp. - with the presence of a conserved 1 nucleotide deletion among members of this group - lemurs (Lemuroidea) and echidna (*Tachyglossus aculeatus*) (Additional Table 11 and File 4). In the case of *Pyroglutamylated RFamide Peptide Receptor* (*QRFPR*), 64 species presented signs of loss of the in-study gene (Additional Table 12), comprising mostly cetaceans and chiropterans (Figures 2 and 3). Accordingly, in all analysed cetaceans the orthologous exon 2 of *QRFPR* was not found and the exon 5 was either not found or exhibited a conserved in-frame premature stop codon (Additional Table 13). Moreover, flying foxes showed, once more, evidence of inactivation with several conserved mutations in their coding reading frame (Additional Table 13). Other examples of species presenting a sequence-altered *QRFPR* includes the California sea lion (*Zalophus californianus*) and echidna (Figure 3; Additional Table 13 and File 5).

**Figure 3:**
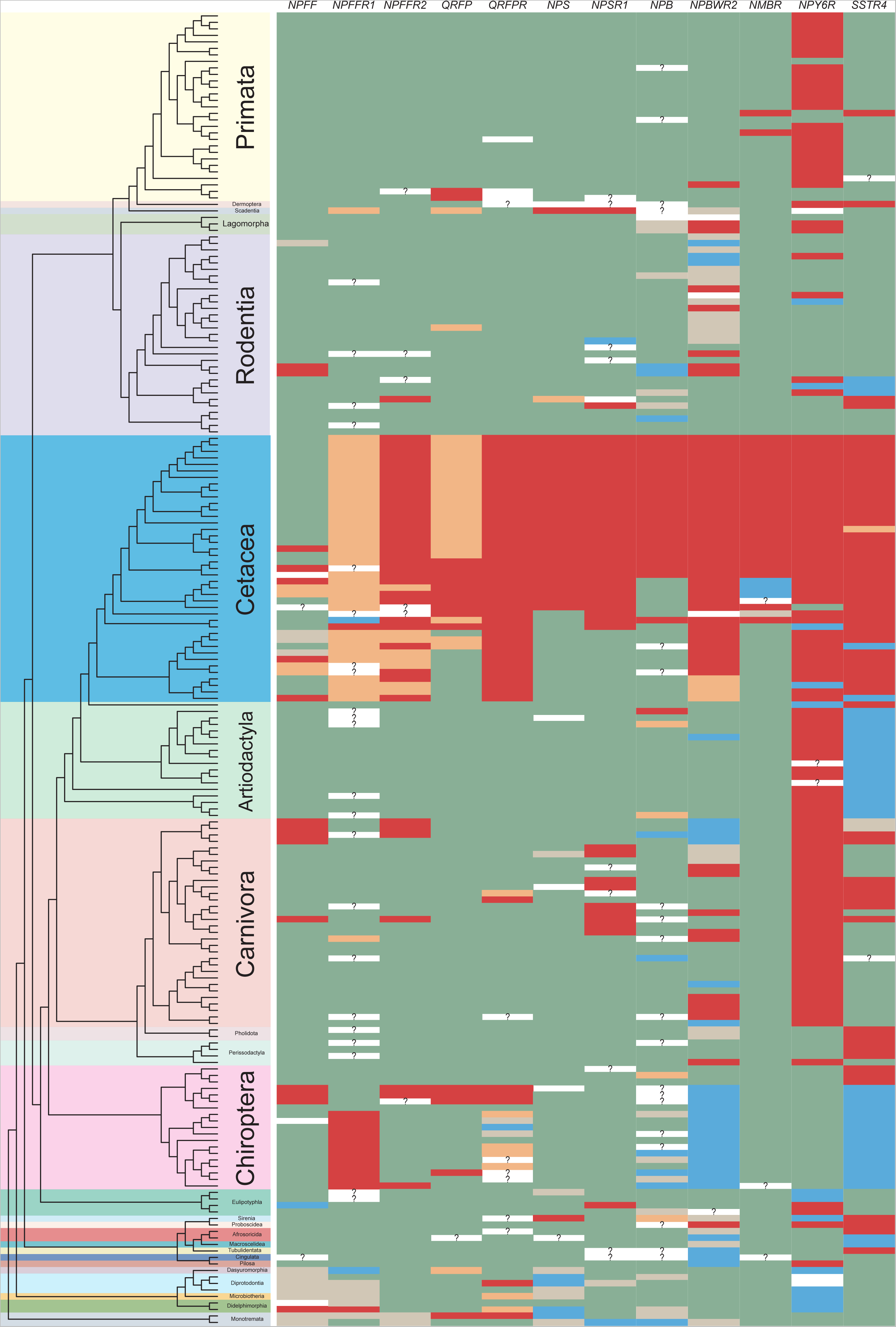
Schematic representation of gene loss events along the mammalian tree. Phylogenetic relationships were adapted from Vazquez [119]. For each gene, we represented in green the species where no ORF-disrupting mutations (frameshift mutations, in-frame premature stop codons, loss of canonical splicing site) were found. On the other hand, species presenting ORF-disrupting mutations are highlighted in red. In orange are symbolised cases where there is some mutational evidence, however the coding status cannot be entirely deduced (e.g., inactivating mutations that occur at the beginning or end of a gene). True absence of exons in the study genes is illustrated in blue. Species presenting (at least) one exon with poor alignment identity with the corresponding reference ortholog are represented in grey. Finally, question marks represent the cases where the absence of a given exon either likely results from the fragmentation (Ns) or incompleteness of the scaffold in the genomic region containing the target gene.

### Convergent gene loss among marine mammals of the Neuropeptide S and receptor

We next examined the coding status of *Neuropeptide S* (*NPS*) and *Neuropeptide S Receptor 1* (*NPSR1*) in mammals to investigate independent gene loss events across the mammalian phylogeny. Although 96 species presented a PseudoIndex higher than 2 for *NPS* (Additional Table 14), only Odontocetes (toothed cetaceans) except sperm whale (*Physeter macrocephalus*) and Indus River dolphin (*Platanista minor*), and West Indian manatee (*Trichechus manatus latirostris*) exhibited valid truncating mutations (Figure 3; Additional Table 15 and File 6). Regarding *NPSR1*, Pseudo*Checker* analyses reported a total of 85 species labelled with a PseudoIndex higher than 2 (Additional Table 16). Annotation of collected genomic sequences revealed *NPSR1* gene erosion mainly in two groups: in toothed cetaceans and aquatic carnivores (pinnipeds and otters) (Figures 2 and 3). While in the first group pseudogenization was confirmed either by the absence of several exons and/or the presence of conserved deleterious mutations, for the latter different ORF-disrupting mutations were retrieved across species (Additional Table 17). In addition to these two groups, we also found evidence of *NPSR1* inactivation in Spanish mole (*Talpa occidentalis*), Damaraland mole-rat (*Fukomys damarensis*) and Chinese tree shrew (*Tupaia chinensis*) (Figure 3; Additional Table 17 and File 7).

### Inactivation of *NPB* and *NPBWR2* genes is found in several mammalian lineages

Sequence search and analysis for *Neuropeptide B* (*NPB*) in 202 mammalian genomes returned a total of 46 species with PseudoIndex higher than 2 (Additional Table 18), the majority due to fragmentation of the genomic region (presence of Ns), true absence of exons, poor alignment identity or incompleteness of the scaffold in the *NPB* genomic region (Additional Table 19). From these species, members of Delphinoidea presented conserved (and validated) frameshift mutations in exon 1 (Figure 2; Additional Table 19 and File 8). With respect to *Neuropeptides B and W Receptor 2* (*NPBWR2*), 107 species displayed a PseudoIndex higher than 2 (Additional Table 20). However, in 52 species with *NPBWR2* putatively pseudogenized, such coding status was predicted due to a true absence of exons, poor alignment identity or fragmentation of the genomic region (presence of Ns) (Figure 3; Additional Table 21). Notwithstanding, ORF-disrupting mutations were identified in 58 species mostly affecting Cetacea and Carnivora orders, but also members of Lagomorpha, Perissodactyla, Proboscidea, Primates and Rodentia orders (Figure 3; Additional Table 21 and File 9).

### Recurrent erosion of a Neuromedin, a Somastotatin and a Neuropeptide Y Receptors in Mammals

We additionally investigated the coding condition of three neuropeptide receptors given their “*Low-quality*” tag in Cetacea genome annotations: *Neuromedin B Receptor* (*NMBR*), *Somatostatin Receptor 4* (*SSTR4*) and *Neuropeptide Y Receptor Y6* (*Npy6r*). For *NMBR*, 35 species displayed a PseudoIndex higher than 2 (Additional Table 22); however, validated ORF-disrupting mutations were only determined in toothed whales and golden snub-nosed monkey (*Rhinopithecus roxellana*) (Figure 3; Additional Table 23 and File 10). On the other hand, a detailed analysis of *SSTR4* occurrence in mammalian genomes revealed an extensive level of gene loss in mammals, with 104 species presenting a PseudoIndex higher than 2 (Additional Table 24). In general, no ORF was detected in the genomes of cetaceans, even-toed ungulates, marine and dog-like carnivorans, bats, pangolins, horses, mole-rats, manatee and aardvark (Figure 3; Additional Table 25 and File 11). For *Npy6r*, a total of 133 species exhibited a PseudoIndex higher than 2 (Additional Table 26), with gene lesion events found and validated in most of cetaceans, even-toed ungulates, primates, carnivores, eulipotyphlans and finally in some rodents (Figure 3; Additional Table 27 and File 12).

### RNA-seq expression

Further examination of several transcriptome datasets of four different cetacean species (Additional Table 2), allowed the assessment of the functional condition of several neuropeptides and receptors reported in this study. Priorly, assessment of the quality of the transcriptomic datasets was performed by checking transcript abundance in genes reported as being highly expressed in a specific tissue, using the Human Protein Atlas has a proxy (http://www.proteinatlas.org). Overall, we could confirm high levels of transcript abundance in the expected tissues, except in testes and brain where no/low expression was found in the expected genes (Additional Table 28).

Across all species, there was a low number of mRNA reads of our target genes, except in *NPFF* (for all species), *NPB* (for *Balaenoptera acutorostrata*) *QRFPR* (for *Balaena mysticetus*) and *NPFFR1*/*NPFFR2* (for *Monodon monoceros*) (Additional Table 29).

After counting the number of spliced reads, exon-intron reads and exonic reads in genes presenting some gene expression, transcriptomic data presented a substantially high proportion of exon-intron reads versus spliced reads in *NPFF*, in stark contrast to the pattern found for *QRFPR* (for *B. mysticetus*), *NPFFR1* (for *Delphinapterus leucas*) and *NPFFR2* (for *M. monoceros*) (Additional Table 29). Moreover, further verification of the presence of ORF disruptive mutations in the produced transcripts, allowed the detection of at least one premature stop codon (or lack of terminal stop codon as observed in *NPB* and *NPFFR1* for Monodontidae) in the transcripts of the analysed cetacean species (Additional Files 13 to 19) revealing either the production of truncated or extended proteins. This is especially relevant in cases where we observed a considerably distinct ratio of exon-intron reads/spliced reads among the remaining species (*QRFPR* for *B. mysticetus, NPFFR1* for *D. leucas* and *NPFFR2 for M. monoceros*) (Additional Table 29).

### Selection analyses indicates relaxation of purifying selection in *NMBR* of Mysticeti

To examine whether genes that are inactivated in odontocetes but not in mysticetes (*QRFP*, *NPS*, *NPSR1*, *NPB* and *NMBR*) exhibit a relaxation in the intensity of selection (both positive and negative), we employed the RELAX method, which explicitly incorporates a selection intensity parameter (k). With the exception of *NMBR*, for which the RELAX test for selection relaxation (k = 0.17) yielded significance (p = 0.01, LR = 10.38), we did not find any evidence for relaxation in the selection intensity for the other genes (Additional Table S30). Further analyses using aBSREL and BUSTED did not reveal any evidence for positive selection in the Mysticeti clade for the *NMBR* gene (Additional Table 31). Therefore, the observed rate of acceleration in mysticetes likely indicates a trend of relaxed purifying selection rather than an episode of positive selection.

## Discussion

From an evolutionary perspective, how the neuronal control of physiological systems modifies and adjusts to radical habitat transitions is mostly unknown. Importantly, these modifications translate into considerable alterations at the anatomical (i.e., pineal gland loss [45]), physiological (i.e., hypoxia-induced adaptations [46]), and behavioural levels (i.e., sleep [47]). Still, the molecular basis of such changes - in perception, motor control, sleep, spatial control, motivation and learning/memory - are challenging to assess. Here, we used as proxy the plethora of neuropeptides known to regulate a vast network of functions: including sleep, feeding, pain, stress, immunity (Figure 1). Some of the reported receptors have previously been identified as lost in early vertebrate lineages, such as *SSTR4* in ray-finned fish [48], or in the ancestors of mammals and birds, like *QRFPR* [49]. Additionally, certain receptors, such as *NPFFR2* [50], *QRFPR* [5], *SSTR4* [48] and *Npy6r* [51], have been reported as lost in other mammals.

Comparative analysis of 40 cetacean genomes uncovered an extensive loss pattern of several neuropeptides and receptors (Figure 2 and 3). The marked gene loss pattern of pleiotropic actors (e.g., [52]), acting on different tissues, anticipates the drastic modification of distinct gene regulatory networks as a result of adaptation to a fully aquatic lifestyle. However, regarding some neuropeptides and receptors (e.g., *NPB*), Mysticeti (baleen whales) seem to retain functional genes (Figure 2). A similar outcome was previously observed in other organ systems (e.g., [53]), thus implying the existence of critical changes in selective pressures over the time in these two related lineages. Nevertheless, our results also suggest that the intensity of purifying selection acting on Mysticeti *NMBR* has been relaxed, which might indicate earlier stages of gene inactivation. Therefore, at least for *NMBR*, it is possible that loss of function has occurred on the stem cetacean branch, rather than on the branch leading to the radiation of odontocetes.

Circadian rhythmicity is one of the most prominent behavioural adaptations in Cetacea. A strong association between the loss of the full melatonin-related gene hub, and the existence of unique bio-rhythmicity in cetaceans has been described [5, 7, 9]. Moreover, disruption of neuropeptides and receptors, such as the neuropeptide Cortistatin and the Dopamine Receptor D5 (*DRD5*), has been correlated with changes in daily activity patterns and energy metabolism [8, 54]. Thus, the absence of a subset of neuropeptides and receptors may have consequences in terms of the neuroendocrine regulation of Cetacea circadian rhythmicity and sleeping behaviour. In agreement, some of the analysed receptors are expressed in hypocretin/orexin neurons (i.e., *NPFFR*s) [55] and *QRFP* was suggested to partially co-localize with orexin [56]. The orexinergic system plays a crucial role in regulating sleep/wake rhythms and thermoregulation, but also in cardiovascular responses, feeding behaviour, spontaneous physical activity and control of energy metabolism [57]. Given the differences in terms of orexinergic bouton density [10] and number of hypothalamic orexinergic neurons [58] found in Cetacea in comparison with their closest relatives (even-toed ungulates), it is expected that disruption of these endocrine messengers may influence cetacean’s unique behaviour. Strikingly, we report the loss of sleep inducing (*QRFP*, *NPB*) [17, 19, 22] or arousal (*NPS*) [24, 26] neuropeptides, and related receptors, in most Cetacea. Yet, sleep-modulators somatostatin (*SST*) and neuropeptide Y (*NPY*) show no signs of erosion [59, 60]. In Cetacea, the loss of multiple neuropeptide sleep regulation seems to have paired with a novel rearrangement of sleep: with the loss of REM sleep and the transition from bihemispheric into unihemispheric sleep [47]. Such transition likely entailed novel regulatory networks acting on a single brain hemisphere rather than the brain as a whole. In agreement, the emergence of Cetacea unihemispheric sleep was accompanied by local thermoregulatory adjustments: uncoupling the brain hemispheric temperature and stabilizing brainstem temperature to prevent sleep-inducing cooling and maintaining motor and sensorial functions (Figure 4) [61, 62]. In addition, hemisphere communication was also adjusted, with the occurrence of interhemispheric fascicle projections, directly into dorsal thalamus nuclei instead of the brainstem projections observed in other mammals [47]. Such functional and anatomical changes possibly allowed the simultaneous vigilance and sleep needed for a mammal to inhabit the high sea.

**Figure 4:**
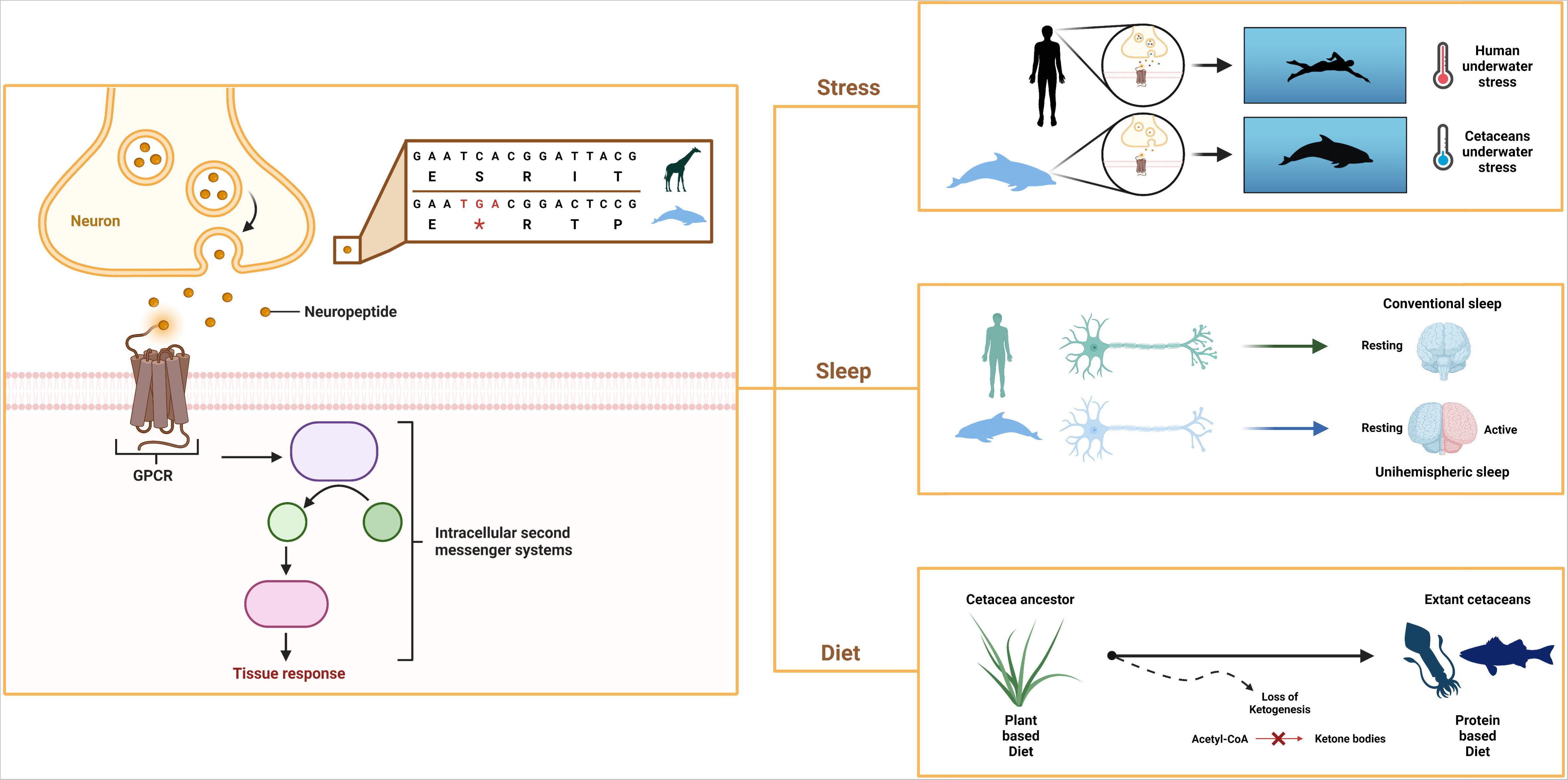
Loss of neuropeptides and receptors contributed to adaptation in an aquatic environment. Impairment of neuropeptides and receptors led to drastic modifications in terms of response to physiological stress, biological rhythmicity and metabolism.

A signalling shift seems to have also occurred in the context of the Cetacea diving response: during which the heart rate is reduced (bradycardia) and the consequent drop in arterial blood pressure is restrained through a strategy of peripheral vasoconstriction, to maintain blood distribution to heart and brain during diving apnoea [63]. Previous reports associated events of gene loss and adaptive gene evolution (positive selection) to the modulation of Cetacea blood volume, pressure and peripheral vasoconstriction [54, 64, 65]. The loss of the hypertensive and heart rate-modulating neuropeptides and receptors (*NPFF*, *QRFP*, *NPB*; cognate receptors and *NMBR*) seems to further contribute to the blood pressure and heart rate regulation [16, 17, 66, 67, 68, 69]. *NPFF* in particular, was shown to be responsible, in rat (*Rattus norvegicus*), for up to 50% of the cardiac component of the baroreflex, participating in the maintenance of the blood pressure levels; and, to modulate the chemoreflex through action on *ASIC* channels, which elicits hyperventilation [16, 66, 70, 71, 72, 73]. *SSTR4* was also suggested to modulate ventilation, with airway contraction observed in knock-out mice [74]. On the other hand, the intact neuropeptides *SST* and *NPY* were suggested to modulate breathing, notably under hypoxic conditions, with *SST* attenuating ventilation and inducing apnoea responses [75, 76]. *NPY* is also known for its prominent effects on the cardiovascular system [77], thus conservation of *NPY* may importantly contribute to blood pressure regulation in this group.

Metabolism is yet another major target of neuropeptide regulation through feeding behaviour and energy expenditure (*NPFF*; *NPB*/*NPW; QRFP; NMBR*) [17, 20, 67, 78, 79, 80] or by controlling insulin secretion (*QRFP*; *NPB*/*NPW*) [18, 81, 82]. Accordingly, cetaceans are known to present unique metabolic needs to adapt to a low glucose diet and to the metabolic constraints of oxygen restriction (Figure 4) [83, 84]. Thus, loss of these feeding suppression neuropeptides and metabolic modulators could contribute to fat signalling and deposition, as well as to the peripheral insulin resistance and high circulating glucose levels under fasting, as observed in odontocete cetaceans [85, 86]. These conditions seem to ensure a steady glucose supply to the central nervous system in the absence of alternative fuel sources - i.e., ketogenesis impairment resulting from genomic loss of hydroxymethyl-CoA synthase - in the context of their low carbohydrate, high protein and fat diet (Figure 4) [87, 88, 89]. In agreement, *NPY*, an inducer of obesity and fat storage was found intact, possibly contributing to the overall fat profile in Cetacea [90]. Although, fat deposition and obesity trigger adipose tissue inflammation in most mammals, Cetacea exhibit a healthy fat phenotype [83, 91]. This could be concurrent with the loss of inflammatory modulation by *NPFF*, *NPS*, or *NPB*/*NPW* [15, 92, 93]. In addition to adipose tissue inflammation, the loss of these neuropeptides and receptors, notably the *SST* receptor *SSTR4* or *NPBWR2* [23, 74], might have further helped cetaceans extend their diving capacities by attenuating lung pain and inflammation, resulting from lung collapse as well as the formation of gas bubbles during deeper dives [65, 93, 94, 95, 96].

Interestingly, we observed a lack of conserved founding mutations common to all analysed Cetacea in most study genes, thus indicating that pseudogenization events took place after the diversification of modern Cetacea lineages (Figure 2). Given the role of neuropeptides as master regulators of a different set of physiological processes, it is expected that, in the course of evolution, molecules with a pleiotropic action should suffer a progressive decay, in parallel with the modifications of the physiological compartments in which they are associated. Adaptation to specific ecological niches might have triggered the occurrence of disrupting mutations in different genomic positions/timescales, thus promoting independent pseudogenization events.

Our approach also identified evolutionary convergence in patterns of gene loss across marine mammals, *NPS* and *NPSR1* (Figure 3). Together, these results suggest that these convergent genomic variations, possibly contributed to the transition from land to water in marine mammal lineages. This is a pattern previously reported for other genes [5, 97, 98] and reinforces the idea of gene loss as a remarkable molecular signature of adaptive evolution in habitat shifts. We also have found other convergent patterns of disruption less easy to explain from an evolutionary perspective. As an example, convergent inactivation of the *NPFF* and receptors (*NPFF* and *NPFFR2* as in Cetacea results on *NPFFR1* coding status were not completely clear) and *QRFP* and receptor in flying foxes (*Pteropus* sp.) and Cetaceans (Figure 3) might be a genomic signature of adaptation to novel niches — an evolutionary convergence previously reported for other genes (e.g., [99]).

## Conclusions

Our findings suggest that alteration/loss of neuropeptide and receptors parallel adaptive shifts in behavioural processes such as bio-rhythmicity, diving and feeding behaviour. On another level, the patterns of gene modification observed in this study, reinforce the idea of gene loss as a major evolutionary driver in the land-to-water habitat transition experienced by the Cetacea ancestor.

## Declarations

### Ethics approval and consent to participate

Not applicable.

### Consent for publication

Not applicable.

### Availability of data and materials

All data generated or analysed during this study are included in this published article and its supplementary information files.

### Authors’ contributions

RV, MC, RR and LFCC conceived the project and designed the research. RV, BP and AM performed the research. RV, MC, FA, ISP, RR and LFCC analysed data. RV and MC wrote the manuscript with input from all authors. All authors have read and approved the final version of the manuscript.

## Acknowledgements

We acknowledge the various genome consortiums for sequencing and assembling the genomes.

## Additional Files

**File name: Additional Tables (.xls)**

**Title of the Data: Additional Tables**

**Description of the Data: Additional Table 1.** Screened mammalian species in the present study, including Cetacea (in grey). This includes, for each species, the corresponding common name, order, inspected genome assembly and database. **Additional Table 2.** In-depth description of the transcriptomic NCBI sequence read archive (SRA) projects, scrutinized in the transcriptomic analysis of the 4 represented cetaceans. **Additional Table 3.** Sequences of non-cetacean mammals incorporated in selection analysis. These include previously annotated genomic sequences at NCBI (Prefix NM_ and XM_). **Additional Table 4.** PseudoIndex value calculated for *NPFF* in selected mammalian species. The table also includes the ID of the genomic sequence from where each target species’ input sequence was extracted, further imported into the analysis. Species displaying a PseudoIndex > 2 are highlighted in red. **Additional Table 5.** Identified *NPFF*-disrupting mutations per mammal species and exon. **Additional Table 6.** PseudoIndex value calculated for *NPFFR1* in selected mammalian species. The table also includes the ID of the genomic sequence from where each target species’ input sequence was extracted, further imported into the analysis. Species displaying a PseudoIndex > 2 are highlighted in red. **Additional Table 7.** Identified *NPFFR1*-disrupting mutations per mammal species and exon. **Additional Table 8.** PseudoIndex value calculated for *NPFFR2* in selected mammalian species. The table also includes the ID of the genomic sequence from where each target species’ input sequence was extracted, further imported into the analysis. Species displaying a PseudoIndex > 2 are highlighted in red. **Additional Table 9.** Identified *NPFFR2*-disrupting mutations per mammal species and exon. **Additional Table 10.** PseudoIndex value calculated for *QRFP* in selected mammalian species. The table also includes the ID of the genomic sequence from where each target species’ input sequence was extracted, further imported into the analysis. Species displaying a PseudoIndex > 2 are highlighted in red. **Additional Table 11.** Identified *QRFP*-disrupting mutations per mammal species and exon. **Additional Table 12.** PseudoIndex value calculated for *QRFPR* in selected mammalian species. The table also includes the ID of the genomic sequence from where each target species’ input sequence was extracted, further imported into the analysis. Species displaying a PseudoIndex > 2 are highlighted in red. **Additional Table 13.** Identified *QRFPR*-disrupting mutations per mammal species and exon. **Additional Table 14.** PseudoIndex value calculated for *NPS* in selected mammalian species. The table also includes the ID of the genomic sequence from where each target species’ input sequence was extracted, further imported into the analysis. Species displaying a PseudoIndex > 2 are highlighted in red. **Additional Table 15.** Identified *NPS*-disrupting mutations per mammal species and exon. **Additional Table 16.** PseudoIndex value calculated for *NPSR1* in selected mammalian species. The table also includes the ID of the genomic sequence from where each target species’ input sequence was extracted, further imported into the analysis. Species displaying a PseudoIndex > 2 are highlighted in red. **Additional Table 17.** Identified *NPSR1*-disrupting mutations per mammal species and exon. **Additional Table 18.** PseudoIndex value calculated for *NPB* in selected mammalian species. The table also includes the ID of the genomic sequence from where each target species’ input sequence was extracted, further imported into the analysis. Species displaying a PseudoIndex > 2 are highlighted in red. **Additional Table 19.** Identified *NPB*-disrupting mutations per mammal species and exon. **Additional Table 20.** PseudoIndex value calculated for *NPBWR2* in selected mammalian species. The table also includes the ID of the genomic sequence from where each target species’ input sequence was extracted, further imported into the analysis. Species displaying a PseudoIndex > 2 are highlighted in red. **Additional Table 21.** Identified *NPBWR2*-disrupting mutations per mammal species and exon. **Additional Table 22.** PseudoIndex value calculated for *NMBR* in selected mammalian species. The table also includes the ID of the genomic sequence from where each target species’ input sequence was extracted, further imported into the analysis. Species displaying a PseudoIndex > 2 are highlighted in red. **Additional Table 23.** Identified *NMBR*-disrupting mutations per mammal species and exon. **Additional Table 24.** PseudoIndex value calculated for *SSTR4* in selected mammalian species. The table also includes the ID of the genomic sequence from where each target species’ input sequence was extracted, further imported into the analysis. Species displaying a PseudoIndex > 2 are highlighted in red. **Additional Table 25.** Identified *SSTR4*-disrupting mutations per mammal species and exon. **Additional Table 26.** PseudoIndex value calculated for *Npy6r* in selected mammalian species. The table also includes the ID of the genomic sequence from where each target species’ input sequence was extracted, further imported into the analysis. Species displaying a PseudoIndex > 2 are highlighted in red. **Additional Table 27.** Identified *Npy6r*-disrupting mutations per mammal species and exon. **Additional Table 28.** Gene expression in different tissues for four different cetacean species. **Additional Table 29.** Transcriptomic read count of *NMBR*, *NPB*, *NPFF*, *NPFFR1*, *NPFFR2*, *NPS*, *NPSR1* and *QRFPR* in *Balaena mysticetus* (bowhead whale), *Balaenoptera acutorostrata* (minke whale), *Delphinapterus leucas* (beluga) and *Monodon monoceros* (narwhal). **Additional Table 30**. Tests for selection relaxation on the Mysticeti branch for *NMBR*, *NPSR1*, *NPB*, *QRFP* and *NPS*. Log-Likelihood Values and Parameter Estimates. **Additional Table 31.** Tests for positive selection on the Mysticeti branch for *NMBR*. Log-Likelihood Values and Parameter Estimates.

**File name: Additional File 1 (.pdf)**

**Title of the Data: Additional Data 1**

**Description of the Data: SRA validation of *NPFF* inactivating mutations in mammals**

**File name: Additional File 2 (.pdf)**

**Title of the Data: Additional Data 2**

**Description of the Data: SRA validation of *NPFFR1* inactivating mutations in mammals**

**File name: Additional File 3 (.pdf)**

**Title of the Data: Additional Data 3**

**Description of the Data: SRA validation of *NPFFR2* inactivating mutations in mammals**

**File name: Additional File 4 (.pdf)**

**Title of the Data: Additional Data 4**

**Description of the Data: SRA validation of *QRFP* inactivating mutations in mammals**

**File name: Additional File 5 (.pdf)**

**Title of the Data: Additional Data 5**

**Description of the Data: SRA validation of *QRFPR* inactivating mutations in mammals**

**File name: Additional File 6 (.pdf)**

**Title of the Data: Additional Data 6**

**Description of the Data: SRA validation of *NPS* inactivating mutations in mammals**

**File name: Additional File 7 (.pdf)**

**Title of the Data: Additional Data 7**

**Description of the Data: SRA validation of *NPSR1* inactivating mutations in mammals**

**File name: Additional File 8 (.pdf)**

**Title of the Data: Additional Data 8**

**Description of the Data: SRA validation of *NPB* inactivating mutations in mammals**

**File name: Additional File 9 (.pdf)**

**Title of the Data: Additional Data 9**

**Description of the Data: SRA validation of *NPBWR2* inactivating mutations in mammals**

**File name: Additional File 10 (.pdf)**

**Title of the Data: Additional Data 10**

**Description of the Data: SRA validation of *NMBR* inactivating mutations in mammals**

**File name: Additional File 11 (.pdf)**

**Title of the Data: Additional Data 11**

**Description of the Data: SRA validation of *SSTR4* inactivating mutations in mammals**

**File name: Additional File 12 (.pdf)**

**Title of the Data: Additional Data 12**

**Description of the Data: SRA validation of *Npy6r* inactivating mutations in mammals**

**File name: Additional File 13 (.pdf)**

**Title of the Data: Additional Data 13**

**Description of the Data: SRA validation of inactivating mutations in *NPFF* transcripts of cetaceans**

**File name: Additional File 14 (.pdf)**

**Title of the Data: Additional Data 14**

**Description of the Data: SRA validation of inactivating mutations in *NPFFR1* transcripts of cetaceans**

**File name: Additional File 15 (.pdf)**

**Title of the Data: Additional Data 15**

**Description of the Data: SRA validation of inactivating mutations in *NPFFR2* transcripts of cetaceans**

**File name: Additional File 16 (.pdf)**

**Title of the Data: Additional Data 16**

**Description of the Data: SRA validation of inactivating mutations in *QRFPR* transcripts of cetaceans**

**File name: Additional File 17 (.pdf)**

**Title of the Data: Additional Data 17**

**Description of the Data: SRA validation of inactivating mutations in *NPSR1* transcripts of cetaceans**

**File name: Additional File 18 (.pdf)**

**Title of the Data: Additional Data 18**

**Description of the Data: SRA validation of inactivating mutations in *NPB* transcripts of cetaceans**

**File name: Additional File 19 (.pdf)**

**Title of the Data: Additional Data 19**

**Description of the Data: SRA validation of inactivating mutations in *NMBR* transcripts of cetaceans**

